# MiR-324-5p regulates the structure of dendritic spines and is essential for hippocampal long-term potentiation

**DOI:** 10.1101/2023.07.19.549725

**Authors:** Emma V Parkins, Darrin H Brager, Jeffrey K Rymer, John M Burwinkel, Diego Rojas, Durgesh Tiwari, Yueh-Chiang Hu, Christina Gross

**Author notes:** **Correspondence should be addressed to:** Christina Gross Cincinnati Children’s Hospital Medical Center Division of Neurology Location R, R2.407 3333 Burnet Avenue Cincinnati, Ohio 45229.

## Abstract

MicroRNAs are an emerging class of synaptic regulators. These small noncoding RNAs post-transcriptionally regulate gene expression, thereby altering neuronal pathways and shaping cell-to-cell communication. Their ability to rapidly alter gene expression and target multiple pathways makes them interesting candidates in the study of synaptic plasticity. Here, we demonstrate that the proconvulsive microRNA miR-324-5p regulates excitatory synapse structure and function in the hippocampus of mice. Both *Mir324* knockout (KO) and miR-324-5p antagomir treatment significantly reduce dendritic spine density in the hippocampal CA1 subregion, and *Mir324* KO, but not miR-324-5p antagomir treatment, shift dendritic spine morphology, reducing the proportion of thin, “unstable” spines. Western blot and quantitative Real-Time PCR revealed changes in protein and mRNA levels for potassium channels, cytoskeletal components, and synaptic markers, including MAP2 and Kv4.2, which are essential for long-term potentiation (LTP). In line with these findings, slice electrophysiology revealed that LTP is severely impaired in *Mir324* KO mice, while baseline excitatory activity remains unchanged. Overall, this study demonstrates that miR-324-5p regulates dendritic spine density, morphology, and plasticity in the hippocampus, potentially via multiple cytoskeletal and synaptic modulators.

## Introduction

Neuronal communication is a highly regulated and dynamic process, requiring rapid changes in protein expression to maintain and shape new connections. The vast majority of excitatory synapses in the brain are comprised of dendritic spines, defined as small protrusions along the dendrite where post-synaptic connections form ^1^. Dysregulation of dendritic spines via altering their morphology, density, or composition can greatly impair neuronal communication. In fact, several neurological disorders have defined dendritic spine pathology and associated defects in plasticity ^2–5^.

One emerging class of spine regulators is microRNA (miRNA). MiRNAs are small noncoding RNA molecules that post-transcriptionally regulate gene expression via targeting messenger RNA (mRNA) ^6^. Their ability to rapidly and post-transcriptionally alter gene expression as well as target multiple proteins and pathways makes them intriguing candidates in the study of synaptic regulation. Subcellular expression studies have identified several miRNAs that are localized to dendritic spines ^7, 8^, and a few miRNAs have indeed been suggested to play important roles in synaptic regulation ^9–11^. In most cases, however, the function of miRNAs in the brain has been studied in the context of neurological disorders, showing altered expression or activity of specific miRNAs in, for example, autism and epilepsy ^12, 13^. The details of how neuronal miRNAs affect neuronal morphology and function in the brain under physiological conditions are not well understood.

One example is miR-324-5p, a proconvulsive miRNA shown to control neuronal hyperexcitability in mouse models of epilepsy ^14–16^. Though miR-324-5p is expressed throughout the body, it is much more highly expressed in the brain than in any other tissue, indicating that it plays an important role in gene regulation in the brain (see: https://ccb-web.cs.uni-saarland.de/tissueatlas/) ^17^. Several studies suggest that miR-324-5p, as well as its target Kv4.2, may modulate dendritic spine formation ^11, 14, 18–21^, but the exact roles of miR-324-5p in structural and functional dendritic spine regulation are unknown.

To analyze miR-324-5p’s physiological roles in the brain, we developed a *Mir324* knockout (*Mir324* KO) mouse model and evaluated dendritic spine morphology and function. It is notable that miR-324-5p shares the pre-miRNA structure with miR-324-3p and that both are encoded by the gene *Mir324*, but miR-324-3p is primarily expressed in other tissues and at very low levels in the brain ^17^, indicating that miR-324-5p is the primary microRNA regulator. Multiple studies support a role of miR-324-5p in neuronal regulation ^11, 14–16, 20^. Using multiple models, we found that dendritic spine density was significantly reduced in *Mir324* KO mice, with loss of spine density comparable to that seen in Fragile X Syndrome and other neurological disorders with changes in dendritic spine density and proven functional consequences ^2, 5, 22, 23^. Morphology of dendritic spines in KO mice also shifted, comprising of fewer thin and increased stubby spines. Acute loss of miR-324-5p via miR-324-5p antagomir (antisense) treatment in adult mice revealed a similar, significant reduction in dendritic spine density, suggesting a role of miR-324-5p in the maintenance of dendritic spines. A candidate-based approach showed dysregulation of proteins involved in dendritic spine and cellular morphology, including the dendritic protein MAP2 and the previously identified target of miR-324-5p, Kv4.2. We further demonstrated that *Mir324* loss impaired hippocampal long-term potentiation (LTP), while presynaptic function appeared to be generally intact. Overall, we demonstrate that miR-324-5p is an important synaptic regulator, modulating dendritic spine form and function while also playing an essential role in LTP.

## Results

### *Mir324* KO mice are viable and healthy

To investigate the role of miR-324-5p in neuronal morphology and plasticity, we generated a *Mir324* knockout (KO) mouse line using CRISPR/Cas9. *Mir324* codes for both miR-324-5p and miR-324-3p. Though both miRNAs are present in the central nervous system, miR-324-5p, not miR-324-3p, is primarily expressed in brain tissue (see: https://ccb-web.cs.uni-saarland.de/tissueatlas/) ^17^, suggesting it is the primary microRNA regulator encoded by *Mir324* in the brain. Successful elimination of miR-324-5p (Figure 1A) and miR-324-3p (Figure 1B) expression was verified using qRT-PCR analysis of *Mir324* KO and WT littermate hippocampal lysates (sequences shown in Figure 1D and Supplementary Figure S1). *Mir324* is encoded within the intron of a protein-coding gene, *Acadv1*. Analysis of Acadv1 mRNA expression in WT and KO mice showed no significant changes (Figure 1C), confirming that phenotypes observed in *Mir324* KO mice are due to lack of expression of *Mir324* and not dysregulated transcription of *Acadv1*. No changes in body weight were observed in 3-4 month-old mice, when experiments were conducted (Figure 1E, mean±SD: WT(male): 25.77±2.13, KO(male):25.37±1.32, WT(female):20.73±1.23, KO(female):20.91±1.93). Reproduction followed a Mendelian distribution (Chi-squared analysis of 50 litters from heterozygous breeding pairs, p=0.07), with no difference in the occurrence of males and females (2-Way ANOVA of sex*genotype, p=0.7) (*data not shown*).

**Figure 1:**
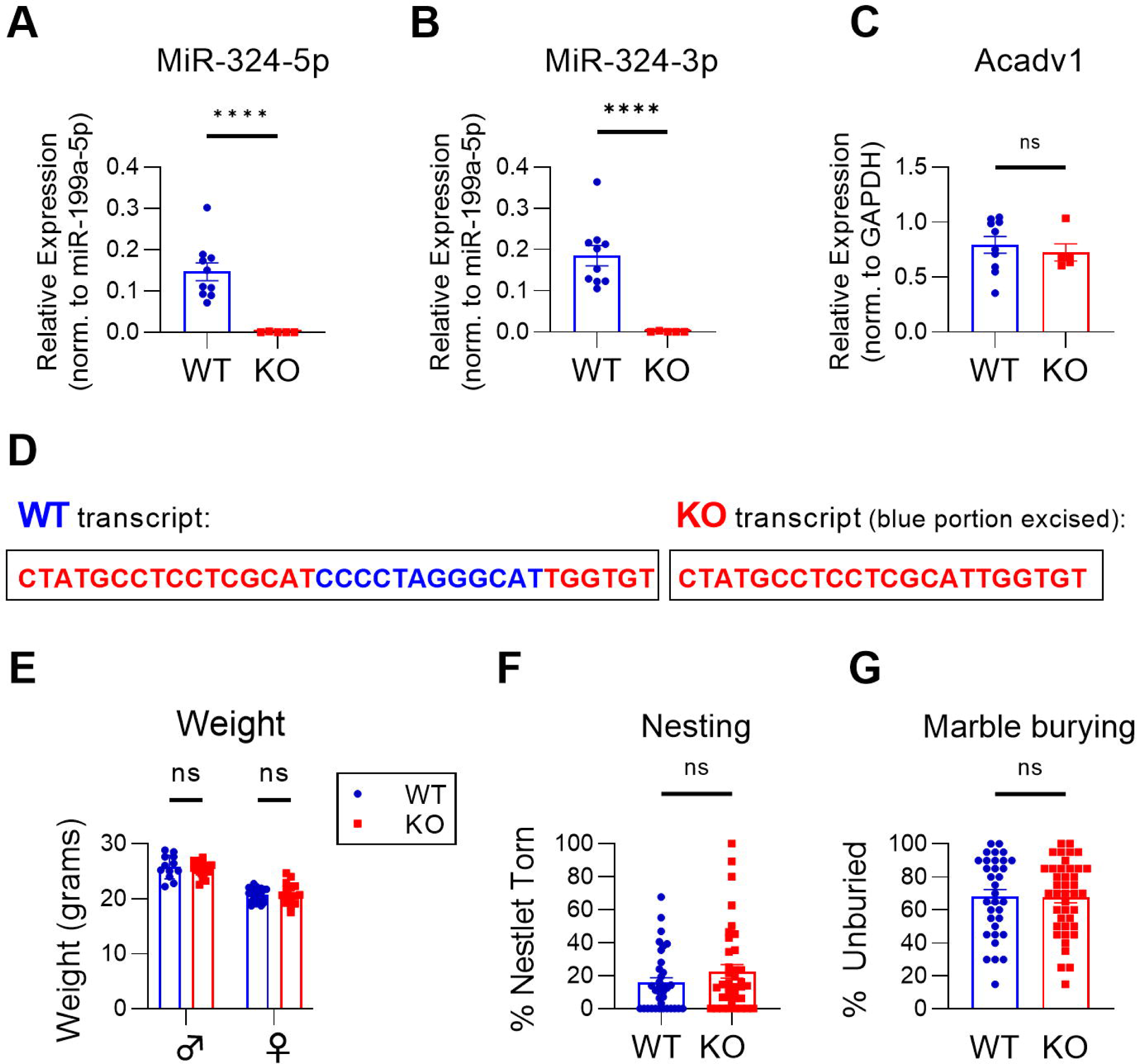
Successful knockout of *Mir324* does not affect host gene expression, weight, or home-cage behavior. (**A-B**) qRT-PCR (quantitative real-time polymerase chain reaction) of *Mir324* KO and WT littermate hippocampal samples confirms loss of miR-324-5p (**A**) and miR-324-3p (**B**) expression in KO mice, confirming successful knockout of *Mir324* (unpaired t-test, n(WT)=10, n(KO)=5; miR-324-5p, p<0.0001; miR-324-3p, p<0.0001). (**C**) *Mir324* is located within an intron of protein-coding gene Acadv1. KO of *Mir324* did not alter Acadv1 mRNA expression (unpaired t-test, n(WT)=10, n(KO)=5; p=0.59.) (**D**) WT and KO transcript sequences. (**E**) Weight varies by sex but not genotype in adult (>2 month old) *Mir324* KO and WT mice (2-Way ANOVA of sex*genotype, n(WT, male)=12, n(WT, female)=23, n(KO, male)=18, n(KO, female)=16; p(interaction)=0.46, p(sex)<0.0001, p(genotype)=0.79). (**F**) Nesting behavior, measured as the percent of nestlet torn after 2 hours, is unaffected in KO mice (Welch’s t-test, n(WT)=35, n(KO)=38; p=0.18). (**G**) Marble burying, quantified by the number of marbles left unburied after 15 minutes, is unaffected in *Mir324* KO mice (unpaired t-test, n(WT)=33, n(KO)=41; p=0.93). Error bars are SEM.

Though *Mir324* KO did not lead to any apparent physiognomic changes, some basic behavioral and morphological assessments were performed to begin characterizing this mouse model. A nesting assay can be used as an indicator of overall health and wellbeing ^24^. No changes in nesting behavior, assessed as the percent of nesting material shredded after 2 hours, were observed (Figure 1F). Marble burying can be used to assess repetitive behavior, which is a feature of autism spectrum disorders (ASD) often used as autistic-like phenotype in mouse models of autism ^25^. No changes in marble burying, assessed as the number of marbles left unburied after 15 minutes, were observed (Figure 1G), indicating that *Mir324* KO mice do not display repetitive behavior. Notably, KO mice did display decreased latency to dig in the marble burying task, a result that may indicate changes in anxiety (mean±SEM in seconds: WT=220±24.3, KO: 166.3±13.56, student’s t-test, n(WT)=39 mice, n(KO)=47, p=0.044).

### *Mir324* KO decreases dendritic spine density and alters dendritic spine morphology

Our previous studies in epilepsy mouse models suggest that miR-324-5p alters the function of excitatory synapse ^14, 15^. We thus analyzed how loss of miR-324-5p expression affects dendritic spine density and morphology in the hippocampus. Global germline loss of *Mir324* reduced dendritic spine density in adult mice as measured in Golgi-stained hippocampus tissue (Figure 2A-B). To analyze dendritic spine morphology in more detail, we utilized *Mir324* KO mice that also carried the Thy1-eGFP transgene to visualize dendrites and dendritic spines in a subset of neurons throughout the CA1 (Figure 2C-E). Analysis of 3-dimensional dendritic spines showed a similar density reduction as observed in Golgi-stained brains (Figure 2C). Assessment of spine morphology (as in ^19, 26^, Figure 2E) also revealed changes in spine morphology, with reduction in the proportion of thin spines and increase in the proportion of stubby spines in KO hippocampus. This reduction in spine density was further supported by reduced PSD95 protein (Figure 2F) and mRNA (Figure 2G) expression. By contrast, synapsin-1 protein expression remained unchanged (Figure 2H). Full blots of cropped images shown in Figure 2F and H are shown in Supplementary Figure S2A and B.

**Figure 2:**
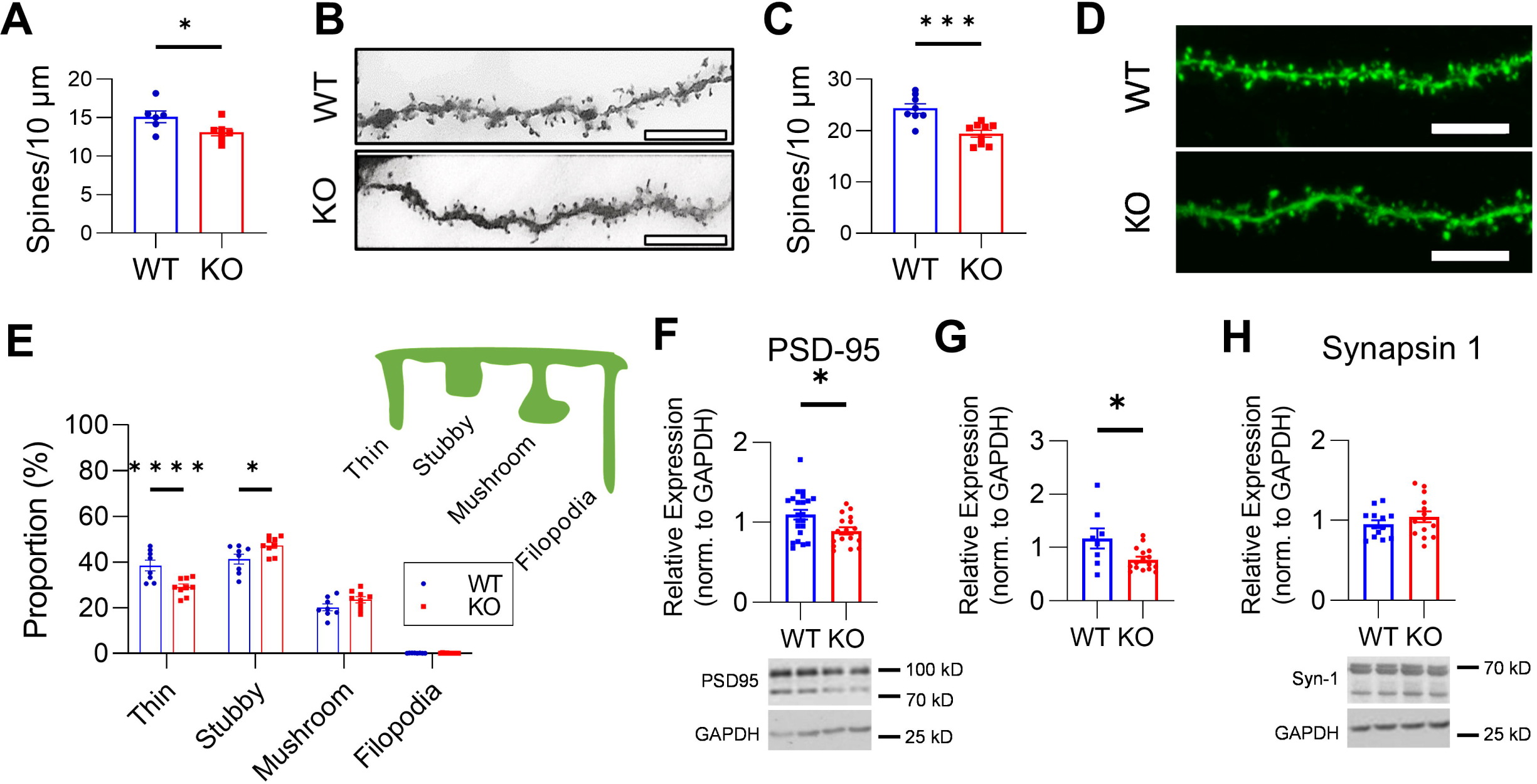
*Mir324* KO decreases dendritic spine density and alters dendritic spine morphology. (**A**) Golgi staining revealed a significant reduction in dendritic spine density in the *Mir324* KO hippocampus (unpaired t-test, n(WT)=6, n(KO)=7, p=0.041). At least 6 segments were counted per mouse across 3 neurons (average 12 dendrites per mouse). (**B**) Sample images of Golgi-stained WT and KO dendrite segments. (**C**) Thy1^hemi^/*Mir324* KO mice also show reduced dendritic spine density in the hippocampus (unpaired t-test, n(WT)=8, n(KO)=9; p=0.0006). Dendritic spines were quantified on secondary apical dendrites of eGFP-positive pyramidal neurons in the hippocampus of PACT-cleared brain sections (4-8/mouse). (**D**) Example images of WT and KO dendrites. Scale bars are 100 μm. (**E**) Spines were then categorized by morphology, revealing a significant reduction in the proportion of thin spines (p<0.0001) and increase in the proportion of stubby spines (p=0.018) in KO dendrites (2-Way RM ANOVA of morphology*genotype with Sidak’s post hoc, p(interaction)<0.0001). Schematic of each dendritic spine morphology category is shown in the upper right corner of panel E. (**F-G**) KO reduces both PSD95 protein (**F**, unpaired student’s t-test, p=0.013, n(WT)=22, n(KO)=17) and mRNA (**G**, unpaired t-test, n(WT)=8, n(KO)=16; p=0.016). (**H**) Synapsin-1 protein expression is unchanged (unpaired t-test, n(WT)=13, n(KO)=14; p=0.29). Scale bars (B, D) are 10 μm. N and dots in diagrams indicate mice. Error bars are SEM. Uncropped blots are shown in Supplementary Figure S2A and B.

### No changes in gross hippocampal or neuronal morphology with *Mir324* deletion

Changes in neuronal and hippocampal morphology are seen in several neurological disorders, including epilepsy ^27^, ASD ^28^, and FXS ^29^. Our previous studies indicate a role of miR-324-5p in epilepsy and seizure susceptibility in a mouse model of temporal lobe epilepsy ^14, 15^. Therefore, to assess if changes in dendritic spine morphology in *Mir324* KO mice are accompanied by potential changes in neuronal morphology in *Mir324* KO mice, we performed Sholl analysis and analyzed the number of intersections (i), and nodes (iii), as well as the length (ii) of dendrites intersecting at each concentric circle of increasing (10 μm) radii (Figure 3B-C). Apical (Figure 3B) and basal (Figure 3C) dendrites were each assessed in addition to total dendrites (*data not shown*). No significant effects of genotype were detected for any of these measures. Likewise, no significant interactions between radii and genotype were detected. Representative WT (blue) and KO (red) neurons are shown in Figure 3A.

**Figure 3.**
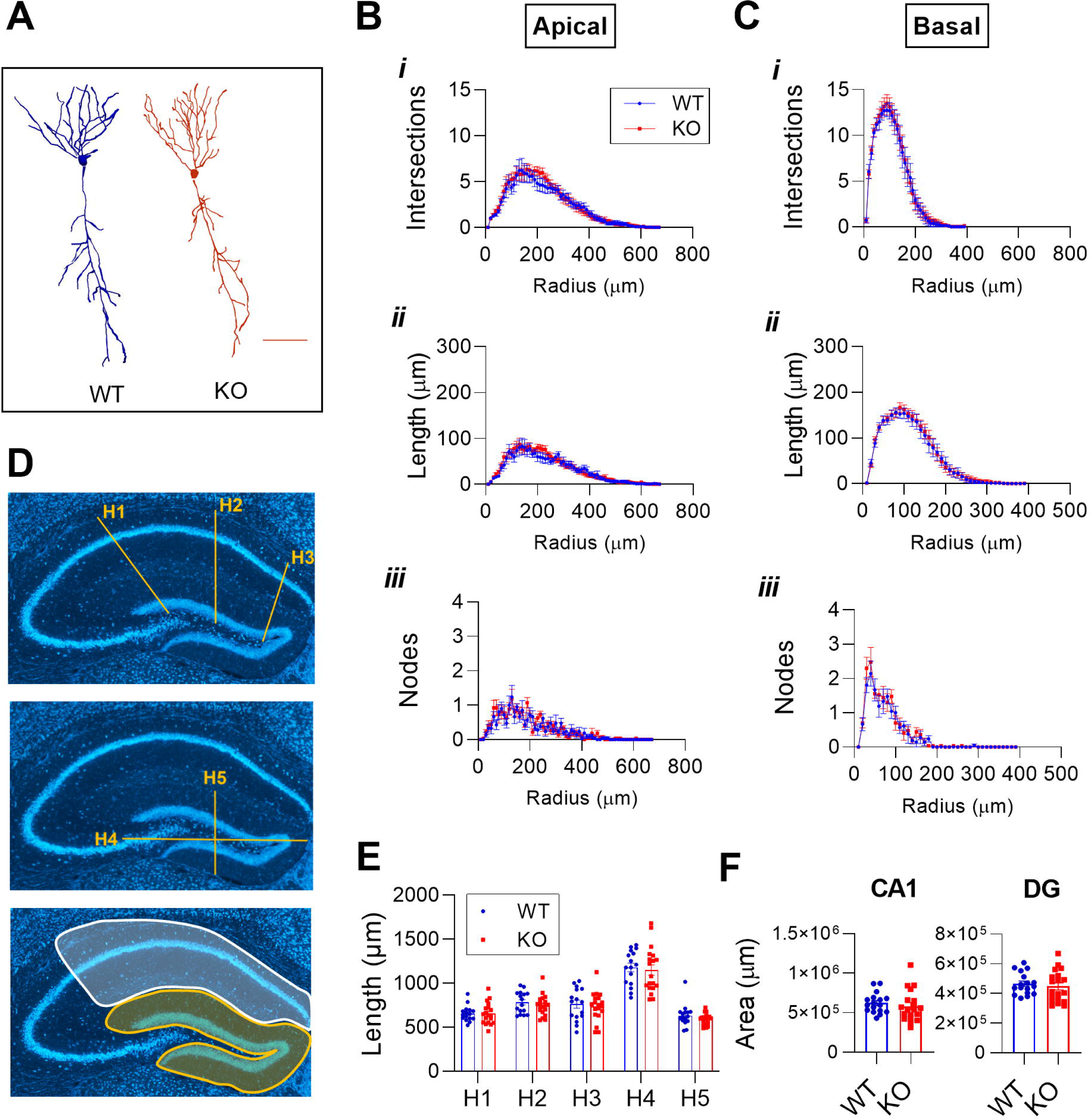
*Mir324* KO does not affect neuronal or hippocampal morphology. Pyramidal neurons in the CA1 subregion of PACT-cleared Thy1^hemi^/*Mir324* KO and WT hippocampi were imaged and traced in Neurolucida (n(WT)=7, n(KO)=9; 3 neurons averaged per mouse). (**A**) Representative images of WT and KO neurons (scale bar is 100 μm). (**B-C**) Sholl analysis revealed no changes in dendritic morphology, with no difference in the number of intersections (€, dendrites that cross over tracings at concentric radii), length (*ii,* sum of the length of dendrites at concentric radii), or nodes (*iii,* branching points identified at concentric radii) (2-Way ANOVA, p(interaction)>0.9 for all measures) in apical (**B**) or basal (**C**) dendrites. (**D**) Representative images of measurements H1-H5 used to assess hippocampal morphology. The top and middle panels illustrate measurements H1-H5, while the bottom panel shows area measurements of the CA1 (in white) and dentate gyrus (DG; in yellow). (**E-F**) No changes in gross hippocampal morphology were observed in *Mir324* KO mice. (**E**) Measurements H1-H5 were unchanged (2-Way ANOVA, n(WT)=16, n(KO)=19; p(interaction)=0.98), and (**F**) both CA1 (unpaired t-test, p=0.48) and DG (unpaired t-test, p=0.56) areas were unchanged. N and dots in diagrams indicate mice. Error bars are SEM.

To assess gross hippocampal morphology, we measured the size and areas of different regions of the hippocampus of *Mir324* KO and WT brain slices stained with Neurotrace (Figure 3D-F, measurements indicated in Figure 3D). No genotype differences in measurements of overall hippocampal size (measurements H1-3, illustrated at the top of Figure 3D) or length and width of the dentate gyrus (measurements H4 and H5, Figure 3D middle) were observed (Figure 3E), and no changes in CA1 or dentate gyrus (DG) area were detected (sample tracings in Figure 3D, bottom; results in Figure 3F). Slices were taken from Bregma levels -1.45 to -1.85 and no differences between genotypes in the bregma levels examined were detected. Our results suggest that loss of *Mir324* specifically regulates dendritic spine morphology without generalized effects at the cellular and subregion scales.

### Acute loss of miR-324-5p reduces dendritic spine density

*Mir324* KO mice show reduced dendritic spine density and altered spine morphology, demonstrating that chronic and complete loss of miR-324-5p alters dendritic spines. Without further study, it is not clear whether this is the result of developmental loss of miR-324-5p, compensation for miR-324-5p loss, or the result of a direct functional relationship. To determine if short-term loss of miR-324-5p had a similar effect on spine density in the CA1 subregion of the hippocampus as KO, adult Thy1-eGFP mice were intracerebroventricularly (ICV) injected with antagomir targeted specifically to miR-324-5p or a scrambled antagomir, and dendritic spine density and morphology were quantified two weeks later (Figure 4A). Just as in KO mice, dendritic spine density was reduced with the miR-324-5p-specific antagomir (Figure 4B). Spine morphology, however, was unaffected (Figure 4C and 4D).

**Figure 4:**
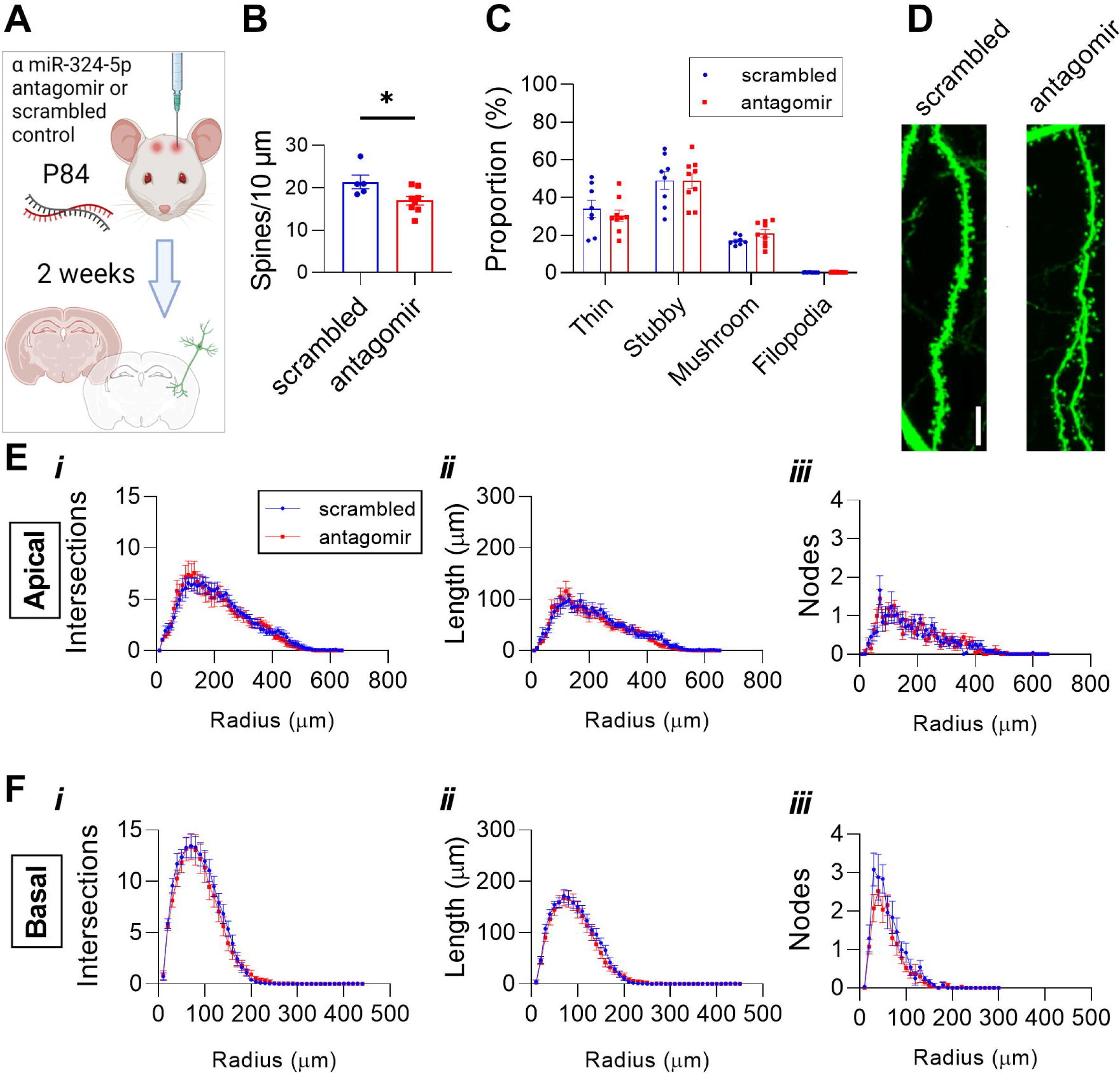
Hippocampal CA1 dendritic spine density is reduced following miR-324-5p antagomir administration. (**A**) Antagomir injection timeline. Briefly, antagomir was ICV injected in P84 Thy1-eGFP^hemi^ mice. Brains were harvested two weeks later and underwent PACT-clearing prior to imaging and analysis. (**B**) Dendritic spine density is reduced with antagomir treatment (student’s t-test, p=0.03; n(scrambled)=6, n(antagomir)=9 mice, 4-6 dendrites per mouse). (**C**) Antagomir treatment does not significantly alter dendritic spine morphology (2-Way ANOVA (category x treatment), p(interaction)=0.1). Representative spine images are shown in (**D**). (**E-F**) Pyramidal neurons in the CA1 subregion of PACT-cleared Thy1-eGFP^hemi^ mice treated with scrambled or miR-324-5p antagomir were imaged and traced in Neurolucida (n(scrambled)=8, n(antagomir)=9; 3 neurons averaged per mouse). Sholl analysis revealed no changes in overall cell morphology, with no difference in the number of intersections (*i*), length (*ii*), or nodes (*iii*) in apical (**E**) or basal (**F**) dendrites (2-Way RM ANOVA; apical: p(interaction)>0.87 and p(treatment)>0.75 for all measures; basal: p(interaction)>0.47 and p(treatment)>0.25 for all measures). Scale bars in (D) are 10 µm. Error bars are SEM.

As in the *Mir324* KO mice, we next analyzed the effects of miR-324-5p inhibition on dendrite morphology. Sholl analysis of miR-324-5p and scrambled antagomir-injected neurons revealed no significant effects of treatment on morphology measured as the number of intersections (i) and nodes (iii), as well as the length (ii) of apical (Figure 4E) and basal (Figure 4F) dendrites at increasing radii of CA1 pyramidal cells relative to scrambled control. Likewise, no significant interactions between radii and antagomir were found. These results suggest that acute miR-324-5p loss-of-function alters dendritic spine density but does not affect dendritic or neuronal morphology in the CA1 subregion.

### Loss of *Mir324* impairs Long-Term Potentiation

To analyze how *Mir324* deletion affects synaptic function and plasticity, we measured excitatory postsynaptic potentials (EPSP) in CA1 pyramidal neurons before and after theta burst pairing (TBP) to induce long-term potentiation (LTP) (Figure 5A). In hippocampal slices from WT mice, TBP produced a significant increase in EPSP slope that persisted for >30 min. By contrast, TBP produced only a transient increase in EPSP slope in hippocampal slices from KO mice that returned to baseline levels (Figure 5B-C). A comparison of EPSPs between baseline and 30 min post-TBP showed that *Mir324* deletion abolished the potentiation in EPSP slope observed in WT hippocampal slices (Figure 5D). Example traces are shown in the bottom panel of Figure 5A. To test if *Mir324* deletion altered baseline release probability, we measured the paired-pulse ratio (PPR) across a range of interstimulus intervals (ISI). We found no difference in the PPR between WT and KO hippocampal slices at any ISI interval (Figure 5E), indicating no change in presynaptic function. Our results show that miR-324-5p expression is essential for LTP.

**Figure 5:**
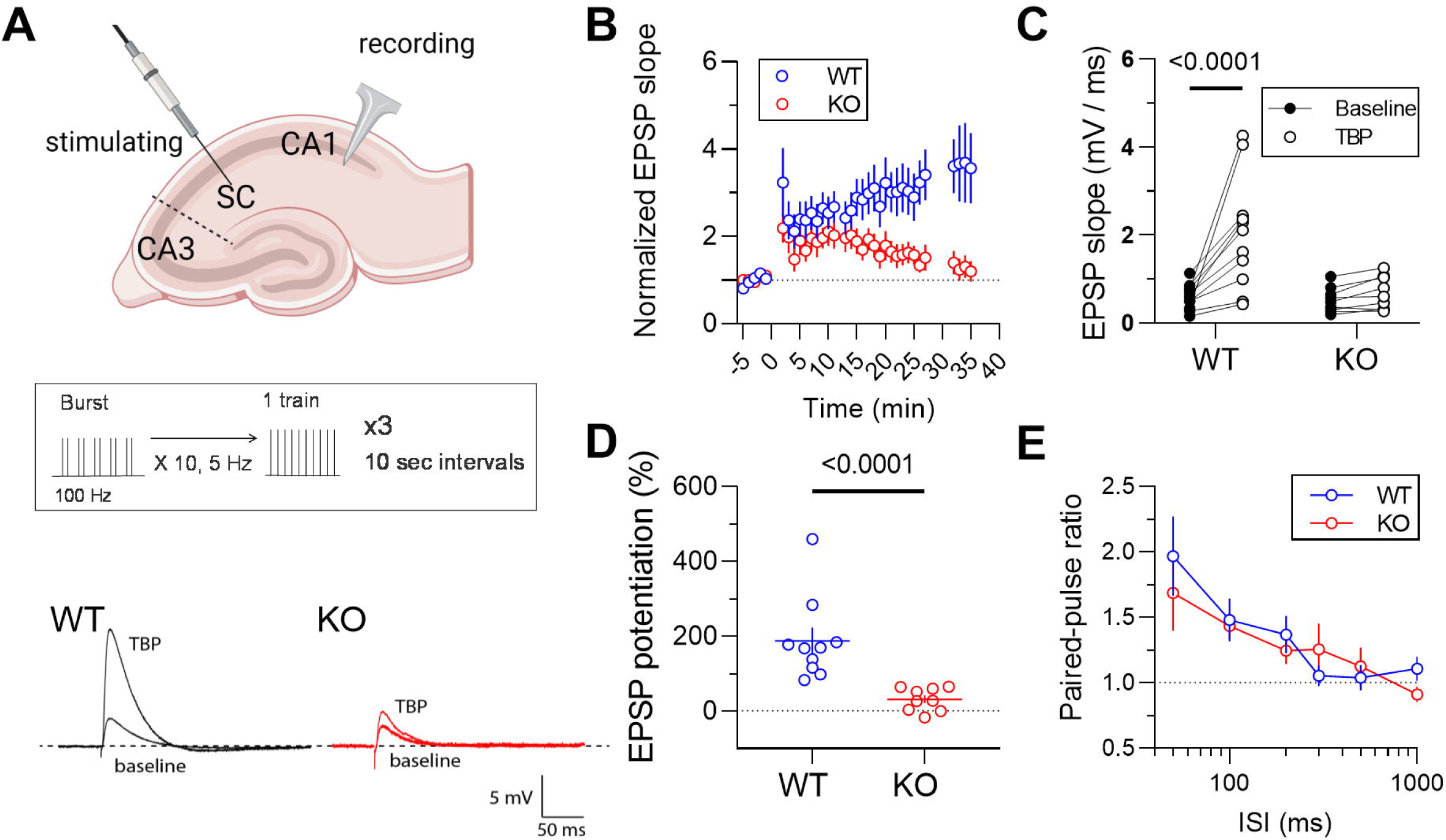
*Mir324* KO impairs LTP but does not affect presynaptic function. (**A**) *Top:* Schematic of LTP induction and recording in hippocampal slices. *Middle:* Schematic of Theta-Burst Pairing protocol. *Bottom:* Example traces of *Mir324* KO (red) and WT (black) neurons at baseline and following theta-burst pairing (TBP). (**B-C**) Excitatory postsynaptic potential slope is significantly increased after TBP in WT but is completely lost in the hippocampi of Mir324 KO mice (**B**, mixed-effects analysis, p(time)<0.0001, p(genotype)=0.049, n(WT)=12, n(KO)=9; **C**, 2-Way ANOVA, p(genotype x pair)=0.004, p(genotype)=0.011, p(pair)=0.0008; Sidak’s posthoc: p(WT-KO baseline)>0.05, p(WT-KO TBP)<0.001. (**D**) Percent potentiation is significantly reduced in KO mice (Mann-Whitney, p<0.0001, n(WT)=10, n(KO)=9. (**E**) Paired-pulse ratio is unchanged with *Mir324* KO (2-Way ANOVA, p(genotype x ISI)=0.66, p(genotype)=0.62, p(ISI)<0.0001, n(WT)=11, n(KO)=10; X is log scale). Error bars are SEM.

### Altered Kv4.2 and cytoskeletal protein expression in *Mir324* KO hippocampus

We next investigated the potential molecular players involved in miR-324-5p mediated dendritic spine regulation. MiR-324-5p targets the mRNA of the voltage-gated A-type potassium channel Kv4.2, and transient inhibition of miR-324-5p using a miR-324-5p antagomir increases Kv4.2 expression in the hippocampus of mice ^14^. Deletion of Kv4.2 leads to enhanced hippocampal LTP ^30^ and heterozygous Kv4.2 loss reduces dendritic spine density ^19^, suggesting that altered Kv4.2 expression could underlie morphological and functional changes in *Mir324* KO neurons. To test if Kv4.2 expression is altered when the *Mir324* gene is deleted permanently, whole hippocampi were dissected from P60 WT and *Mir324* KO mice and Kv4.2 expression was assessed via Western blot. We found that, similar to antagomir treated hippocampi, Kv4.2 protein (Figure 6A) and mRNA expression (Figure 6B) were significantly higher in the KO hippocampus, suggesting that altered Kv4.2 expression could contribute to synaptic dysfunction and altered dendritic spine morphology. By contrast, potassium channels KCNQ2 (Figure 6C) and KCNQ3 (Figure 6D), which do not have direct target sequences for miR-324-5p, were not affected.

**Figure 6:**
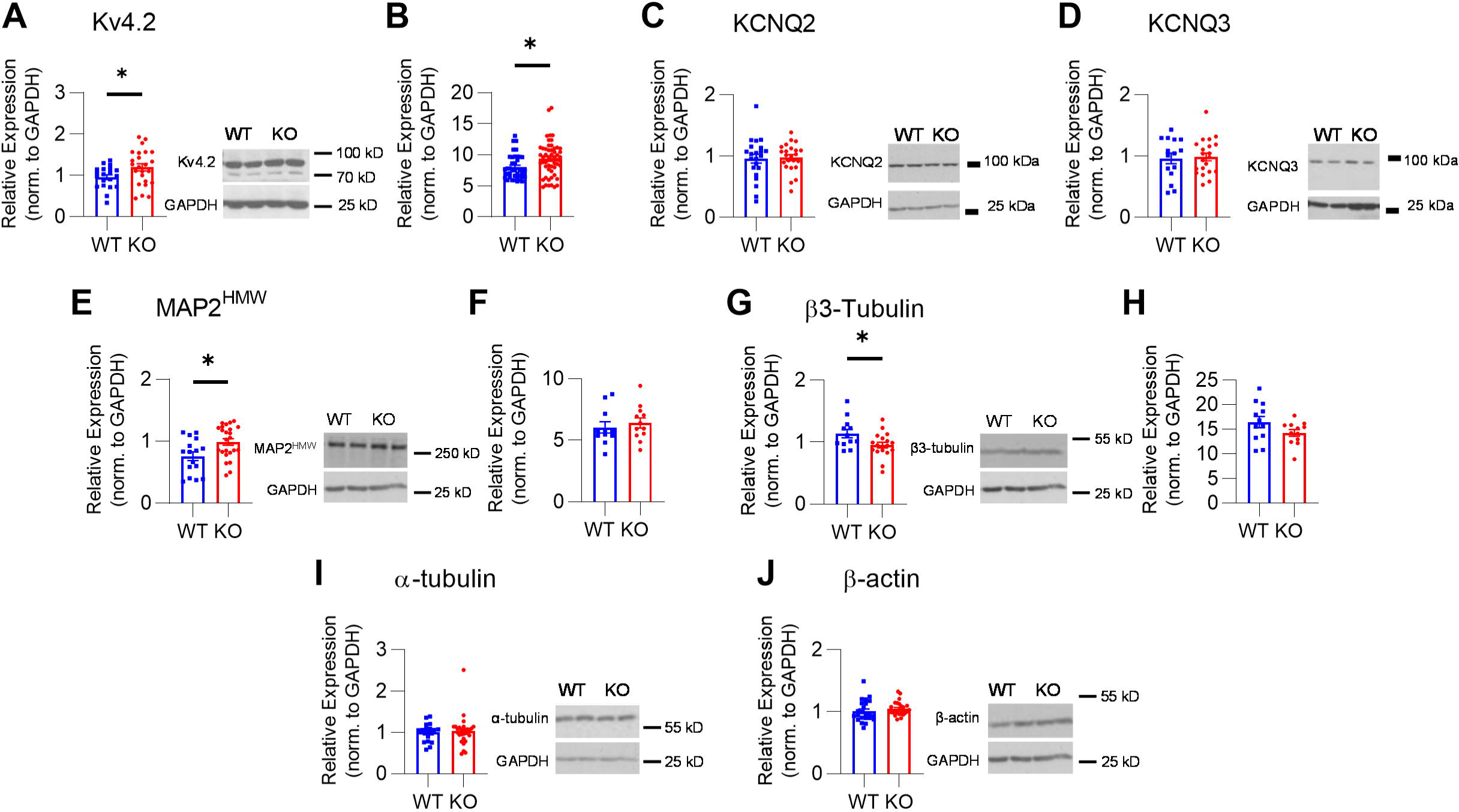
*Mir324* KO alters expression of potassium channels and cytoskeletal proteins in the hippocampus. (**A-B**) KO increases Kv4.2 protein (**A**, unpaired t-test, p=0.038, n(WT)=20, n(KO)=24) and mRNA expression (**B**, unpaired t-test, p=0.015, n(WT)=39, n(KO)=54). (**C**) KCNQ2 expression is not affected by loss of *Mir324* (unpaired t-test, n(KO)=23, n(WT)=22, p=0.814). (**D**) KCNQ3 expression is also not affected (unpaired t-test, n(KO)=21, n(WT)=15, p=0.827). (**E-F**) *Mir324* KO increases MAP2^HMW^ protein (**E**, Welch’s t-test, p=0.015, n(WT)=16, n(KO)=24) but not mRNA (**F**, unpaired t-test, n(WT)=11, n(KO)=12; p=0.56) expression. (**G, H**) β3-Tubulin protein expression is reduced in KO hippocampus (**G**, unpaired student’s t-test, p=0.029, n(WT)=12, n(KO)=16) but mRNA expression is unchanged (**H**, student’s t-test, n(WT)=12, n(KO)=12; p=0.11). (**I-J**) No changes in expression of α-tubulin (**I**; Welch’s t-test, n(WT)=18, n(KO)=27; p=0.75) or β-actin (**J**; Welch’s t-test, n(WT)=23, n(KO)=27; p=0.378), or were detected. Representative blots are shown to the right of protein quantifications. Error bars are SEM. Uncropped blots are shown in Supplementary Figure 2C-I.

MiRNAs target many mRNAs, suggesting that dysregulation of other proteins in addition to Kv4.2 may contribute to impaired neuronal function and altered dendritic spine morphology. Indeed, pilot RNA-Seq studies suggested changes in cytoskeletal proteins in the hippocampus of *Mir324* KO mice (*data not shown*), which could contribute to altered dendritic spine morphology and synaptic plasticity. We thus quantified the expression of select cytoskeletal proteins shown to be key for dendritic spine morphology using Western blotting. Specifically, we measured expression of MAP2 (high molecular weight; MAP2^HMW^), β3-tubulin, α-tubulin, and β-actin proteins. MAP2^HMW^ expression was increased (Figure 6E), while mRNA levels remained the same (Figure 6F). Notably, MAP2^HMW^ plays a critical role in LTP induction; MAP2^HMW^ translocation from the dendritic shaft to spine head is essential for AMPA receptor insertion and spine morphology changes in LTP, and loss impairs LTP induction ^31^. Somewhat in keeping with the observed reduction in spine density, β3-tubulin expression was reduced in KO hippocampi (Figure 6G), while mRNA expression remained unchanged (Figure 6H). *Mir324* KO did not affect expression of α-tubulin or β-actin (Figure 6I-J). Whole uncropped western blots used for example images in Figure 6 are shown in Supplementary Figure S2C-I.

## Discussion

This study shows that miR-324-5p plays an important role in structural and functional dendritic spine regulation. Chronic, complete loss of miR-324-5p via knockout of *Mir324* in mice reduces dendritic spine density (Figure 2A-D) and shifts the morphological composition of dendritic spines, decreasing the proportion of dendritic spines with thin, “immature” ^3, 32–34^ morphology and increasing the proportion of stubby, potentially more stable spines (Figure 2E). Acute loss of miR-324-5p results in a similar reduction in spine density (Figure 4B), indicating that miR-324-5p regulates dendritic spines continuously. Additionally, we found that *Mir324* KO has functional consequences, with KO dramatically impairing hippocampal LTP (Figure 5B-D). As potential underlying molecular contributors we detected changes in expression of cytoskeletal and synaptic proteins, including increased expression of MAP2^HMW^ and Kv4.2 (Figure 6A,E). Both MAP2^HMW^ and Kv4.2 play important roles in LTP ^30, 31^ and regulate dendritic spines ^19, 31, 35^. Future studies will have to explore the role of MAP2, Kv4.2 and other targets in miR-324-5p-mediated regulation of dendritic spines and LTP.

Our study provides insight into the role of miR-324-5p in synaptic regulation, an important step in understanding the role miRNAs play in neuronal communication, synchronization, and activity. *Mir324* KO impairs LTP without affecting presynaptic function (Figure 5), strongly suggesting that this miRNA may regulate synaptic plasticity. These findings are in line with a few other studies that suggest a role of miR-324-5p in synaptic plasticity. MiRNA sequencing of the barrel cortex in a mouse model of associative memory revealed changes in the expression of several miRNAs, including miR-324-5p ^21^. Concomitant antagomir-induced silencing of miR-324-5p and miR-133a demonstrated that loss of both miRNAs at the same time reduces dendritic spine formation in associative memory ^18, 20, 36^, but does not provide evidence for the role of miR-324-5p itself. Taken together with our findings, these studies suggest that miR-324-5p plays a role in learning and memory.

MiRNAs are dynamic molecules, with temporal and spatial-specific expression patterns that can result in different profiles of expression and function in different brain subregions and at different ages ^37, 38^. The same is true of dendritic spines ^33^. Here, we focus on dendritic spine changes in secondary apical dendrites within the CA1 subregion of adult mice. We chose to focus on the CA1 subregion because it is the area of the hippocampus with highest Kv4.2 expression ^39^, which we have shown previously plays an important role in the anticonvulsant effect of miR-324-5p antagomir treatment in mice ^14^. Kv4.2 expression and dendritic spine phenotype both vary by subcellular location, with different rates of Kv4.2 turnover and altered expression patterns at locations proximal and distal to the soma ^40^. We sought to minimize the effects of subcellular localization and cell location within our analysis by including dendrites proximal and distal to the soma and from neurons located across the CA1. Because of this, subcellular-, layer-, and location-specific regulation is lost in our analysis. Further work is needed to determine the relationship between subcellular and subregional location and miR-324-5p-mediated spine regulation.

Our results show a shift towards less thin dendritic spines and more stubby spines in *Mir324* KO mice. These morphological categories of spines are thought to indicate changes in synaptic strength, with thin spines identified as less mature and stable ^3, 32, 33^. In line with this interpretation of the observed phenotype, we show that synaptic function is indeed altered, as illustrated by strongly impaired LTP.

Of note, acute knockdown of miR-324-5p with an antagomir reduced dendritic spine density to a similar extent as observed in *Mir324* KO mice. This suggests that miR-324-5p is important for dendritic spine maintenance in adult mice and may be involved in dendritic spine dynamics, which should be explored in the future. While the reduction in dendritic spine density is relatively small (∼15-20%), the magnitude of changes is similar as in diseases leading to impairments in neuronal function, such as Fragile X Syndrome ^2^, suggesting that these changes have functional consequences as supported by our data (Figure 5) and previous publications ^14, 15^.

Understanding the miRNA-mediated mechanisms underlying regulation of dendritic spine morphology and dynamics is important not only to reveal how miRNAs control neuronal function and plasticity but has potential translational application as well. Though their dendritic spine phenotypes vary, several neurological disorders have deficits in plasticity and defined dendritic spine pathology ^2, 4, 5, 41^. For example, dendritic spines are characteristically changed in epilepsy, where spine alterations may be a cause and consequence of seizure activity ^42^. Previously, our lab has shown that an antagomir to miR-324-5p reduces seizure susceptibility and frequency in mouse models of epilepsy ^14, 15^, which may point towards a novel therapeutic target. In fact, it is known that miRNAs can serve as drug targets, and there are several studies and clinical trials assessing miRNA inhibition as a therapeutic strategy in other diseases ^43, 44^. Thus, beyond the contribution this research makes to our understanding of miRNA in neuronal function, it may also help further develop a miRNA target for this new therapeutic approach.

## Material and Methods

### Animals

All animal procedures were approved by the Institutional Animal Care and Use Committees of CCHMC and UT Austin and complied with the Guideline for the Care and Use of Laboratory Animals, under the Animal Welfare Assurance numbers A3108-01 (CCHMC) and A4107-01 (UT Austin). All methods are reported in accordance with ARRIVE guidelines. Thy1-eGFP^hemi^ mice (RRID:IMSR_JAX:007788) were obtained from Jackson Labs and bred with C57BL/6J wild type mice (RRID:IMSR_JAX:000664) in-house. Three-month-old Thy1-eGFP^hemi^ mice were used for bilateral intracerebroventricular injection of antagomir. *Mir324* KO mice were generated by the CCHMC Transgenic Animal and Genome Editing Core Facility using CRISPR/Cas9 gene editing of C57BL/6N mice and their tissue collected for Golgi staining and protein/mRNA/miRNA quantification at 8-10 weeks of age, and for slice electrophysiology at 2-4 months of age. For fluorescent imaging and analysis of spine morphology, *Mir324* KO mice were bred with Thy1-eGFP^hemi^ mice to generate miR-324 het/Thy1-GFP^hemi^ mice. These mice were mated to generate sex-matched *Mir324* WT/Thy1-GFP^hemi^ (WT) and *Mir324* KO/Thy1-GFP^hemi^ (KO) mice. Pups were weaned at P28 and were housed with same sex littermates (minimum 2 and maximum 4 per cage) in a standard cage with food and water provided ad libitum. A standard mouse house was kept in every cage. Mice were maintained on a standard 14:10 hour light:dark cycle at CCHMC and 12:12 hour light:dark cycle at UT Austin, and all experiments were performed during the light cycle.

### Generation of *Mir324* KO mice

The sequence for mmu-miR-324-5p is 22 nucleotides long (MIMAT0000555_ is 5’ – CGCAUCCCCUAGGGCAUUGGUGU – 3’). The sgRNA target sequence (5’ – GCTTTACACCAATGCCCTAG – 3’) was selected according to the location and the on- and off-target scores from a web tool CRISPOR.org (https://pubmed.ncbi.nlm.nih.gov/27380939/). The full *Mir324* transcript and relevant sites are shown in Supplementary Figure S1. The sgRNA was in vitro synthesized using the MEGAshorscript T7 kit (ThermoFisher, Cat. No. AM1354) and purified by the MEGAclear Kit (ThermoFisher, Cat. No. AM1908), following manufacturer’s instructions. sgRNA and Cas9 protein (ThermoFisher, Cat. No. B25641) were mixed at the final concentration of 75 and 150 ng/ul, respectively, and incubated at 37°C for 15 min to form a ribonucleoprotein complex, which was subsequently injected into the cytoplasm of one-cell-stage embryos of the C57BL/6 background using a piezo-driven microinjection technique as described previously (Scott & Hu, 2019). Injected embryos were immediately transferred into the oviducal ampulla of pseudopregnant CD-1 females. Live born pups were genotyped by Sanger sequencing and confirmed one founder mice, in which a 12 bp portion within the seed region of the *Mir324* gene was deleted. *Mir324* KO mice were crossed with C57BL/6J mice to maintain an active colony. Genotype of the offspring was originally confirmed by Sanger sequencing followed by ear-notch PCR for colony maintenance (forward primer WT: 5’ – CATCCCCTAGGGCATTGGT – 3’; forward primer KO: 5’ – CTATGCCTCCTCGCATTGG – 3’; reverse primer (both): 5’ – GTTTGGGGACAAAATTCACAACT – 3’).

### Nest building assay

Nesting behavior was assessed as previously described ^45^ with mice at 5 weeks of age. Briefly, untorn nestlets (approximately 3 grams) were weighed at 0 and 2 h after addition to a standard mouse cage with a single-housed mouse. WT and KO littermates of both sexes were utilized. Several mice were excluded from analysis due to cage flooding (n(KO)=3).

### Marble burying assay

Marble burying behavior was tested two weeks following assessment of nest building behavior in the same mice. Marble burying was assessed similarly as in ^46^. Briefly, following 1 hour of acclimation to home cages placed under a sterile hood, mice were placed in 10.5×19 inch standard rat cages with twenty blue small glass marbles arranged in a 5 × 4 grid on fresh bedding (ca. 8 cm deep). Latency to start digging was measured as the seconds lapsed before a mouse begins to demonstrate any digging behavior. After 15 minutes marbles covered 50% or more were scored as “buried”. Mice that did not bury any marbles were excluded as “nonparticipants” and not included in analysis (n(WT)=2).

### Golgi stain imaging

Dendritic spines of CA1 pyramidal neurons (bregma level -1.9 to -2.2) were assessed using Rapid Golgistain Kit (FD Neurotechnologies, MD) as described previously^47^. Harvested brains were subjected to Golgi impregnation, sectioned at 120 µm thickness, and stained as per manufacturer’s protocol. Sections were imaged using a Nikon inverted microscope with 4X, 20X, and 60X/NA1.4 oil immersion lenses. Dendritic spines of CA1 pyramidal neurons located across the CA1 were assessed on secondary dendrites (60-120 μm in length) located at least 50 μm from the cell body. On average, 12 dendrites (+/- 1 SEM) from 3 neurons were assessed per mouse by experimenters blinded to genotype and sex. Spines were manually counted on ImageJ (NIH) and spine counts of all neurons from each WT and *Mir324* KO mouse were pooled for statistical analysis.

### Nissl staining and hippocampal measurements

Following deep anesthesia with at least 200 mg/kg pentobarbital, mice were intracardially perfused with 2% paraformaldehyde (PFA). Whole brains were removed and preserved in 4% PFA overnight at 4LJC prior to cryoprotection with sucrose solution (24 hr 10%, 20%, and 30% sucrose at 4LJC) and cryopreservation at -80LJC. Brains were embedded in Tissue-Tek OCT compound and sectioned at 20 µm, then stained with Neurotrace (435/455, Fisher Scientific, N21479) according to the product’s protocol. Slides were imaged and measurements of the hippocampus were obtained via Nikon Elements (Tokyo, Japan, RRID:SCR_014329) and ImageJ (RRID:SCR_003070) software and assessed individually. Measurements are outlined in Figure 2A. Briefly, measurement H4 transects the dentate gyrus from dentate granule cell dendrites at the apex to the end. H5 bisects H4 and extends across the shells of the dentate, from dendrites at the supramedial to infrapyramidal blades. H1-3 evenly transect H4, originating beneath the granule cell layer and ending at the tip of basal dendrites of CA1 pyramidal cells. Brain sections were assessed for each mouse at the approximate bregma level of -1.7 mm. Contrast and brightness were adjusted for visualization in Figure 2.

### Thy1-eGFP^hemi^ fluorescent imaging

Thy1-eGFP^hemi^ mice and *Mir324* KO mice were crossbred to generate Thy1-eGFP^hemi^/*Mir324* KO mice. Brains from sex- and age-matched Thy1-eGFP^hemi^/*Mir324* KO and WT mice were collected on postnatal day 60 (P60), fixed with 2% PFA, and PACT-cleared as previously described ^19^. Refractory Index-Matching Solution (RIMS)-mounted brains were imaged on Nikon A1R LUNV with 4X, 20X, and 60X water immersion lenses. Shot noise was removed from images using the Nikon NIS Denoise.ai (Nikon Instruments Inc., RRID:SCR_014329). Six to eight secondary dendrites across 3-4 pyramidal neurons in the CA1 were imaged per mouse and dendrites between 60 and 120 μm in length were included in analysis. Dendritic spines of CA1 pyramidal neurons located across the CA1 were assessed on secondary dendrites located at least 50 μm from the cell body. Neuronal morphology, dendritic spine density, and spine morphology were assessed using Neurolucida 360 (RRID:SCR_016788) and Neurolucida Explorer (MBF Biosciences, RRID:SCR_017348) with default settings as described ^26^. Spine morphology was classified using a categorization “macro” in the Neurolucida software, and based on classifications outlined by Rodriguez et al., 2008 ^48^ as pre-set by the software. Briefly, dendritic spines were classified based on their head-to-neck and length-to-head ratios, as well as head width and overall length. Thin spines and filopodia were identified as having a head-to-neck ratio of smaller than 1.1, encompassing spines with and without small heads, and were distinguished by length, with filopodia being longer than 3μm ^48^.

### Antagomir injection surgery

Three-month-old Thy1-eGFP^hemi^ mice (weight (mean±SD): Male: 25.57±2.17, Female: 19.43±0.51) received 0.5 nmol (in 2μl) of either scrambled or miR-324-5p specific antagomirs in artificial cerebrospinal fluid (aCSF) via intracerebroventricular injection, as in ^14^. Mice were assigned to groups in a randomized way, and where possible, equal numbers of mice from one litter were injected with miR-324-5p-specific antagomir and scrambled antagomir. Briefly, mice were anesthetized with 4% isoflurane. Mice remained under light isoflurane anesthesia (approximately 1%), and respiration patterns were monitored throughout. Carprofen (5 mg/kg) was administered subcutaneously and allowed to absorb completely (3-5 minutes) both before and after surgery. The head was shaved and disinfected using Dermachlor (2% Chlorhexidine), then the skull was exposed by making an incision along the midline. Dorsoventral coordinates were measured from bregma and two holes were drilled at APLJ=LJ-3.0LJmm; LLJ=LJ± 1.0LJmm and VLJ=LJ2.0LJmm. Antagomir was administered slowly over a 5 min duration using a 5 μl Hamilton syringe, which was left in place for 10 min to allow the diffusion of the injected volume. The needle was retracted slowly over a 5 minute period. Tissue was collected two weeks after antagomir injection.

### Long-Term Potentiation / slice electrophysiology

#### Preparation of acute hippocampal slices

All experiments were conducted in accordance with the University’s Institutional Animal Care and Use Committee. Hippocampal slices (300 µm) were prepared from 2- to 4-month-old mice as described in ^49^. Briefly, animals were deeply anesthetized with a lethal dose of ketamine and xylazine and intracardially perfused with ice-cold modified aCSF containing the following (in mM): 210 sucrose, 2.5 KCl, 1.2 NaH_2_PO_4_, 25 NaHCO_3_, 0.5 CaCl_2_, 7.0 MgCl2, and 7.0 dextrose bubbled with 95% O_2_/5% CO_2_. The brain was removed and bisected along the midline, an oblique cut was made to promote the planar orientation of the dendrites, the brain was mounted to the stage of a Vibratome, and sections were made from the middle hippocampus. Slices were placed in a holding chamber with aCSF containing the following (in mM): 125 NaCl, 2.5 KCl, 1.25 NaH_2_PO_4_, 25 NaHCO_3_, 2.0 CaCl_2_, 2.0 MgCl_2_, and 21 dextrose, pH 7.4, bubbled with 95% O_2_/5% CO_2_ at 35°C for 45– 60 min and then kept at room temperature.

#### Electrophysiology

Slices were placed individually, as needed, into a submerged recording chamber and continuously perfused with oxygenated extracellular saline containing the following (in mM): 125 NaCl, 3 KCl, 1.25 NaH_2_PO_4_, 25 NaHCO_3_, 2.0 CaCl_2_, 1.0 MgCl_2_, and 21 dextrose, pH 7.4, at 32–34°C. Slices were viewed with a Zeiss AxioExaminer D microscope fitted with a 60 water-immersion objective and Dodt contrast optics. Patch pipettes were pulled from borosilicate glass and wrapped with Parafilm to reduce capacitance.

#### Whole-cell recording

For whole-cell recording, pipettes were filled with the following (in mM): 120 K-gluconate, 20 KCl, 10 HEPES, 4 NaCl, 4.0 Mg-ATP, 0.3 Na-GTP, and 14 K2-phosphocreatine, pH 7.3 with KOH. Whole-cell recordings were made using a Dagan BVC-700A in current-clamp mode. Data were sampled at 40 kHz, analog filtered at 5 kHz, and digitized by an ITC-18 interface connected to a computer running Axograph X. Series resistance (*R*S) was monitored throughout the recording and the experiment was discarded if *R*S exceeded 30 MΩ or varied by 20%. GABAA- and GABAB-mediated IPSPs were blocked by 2 µM SR95531 and 5 µM CGP55845, respectively. To prevent epileptiform activity, a cut was made between area CA3 and area CA1. Schaffer collateral EPSPs of 2–3 mV were elicited using tungsten-stimulating electrodes placed 20 µm from the apical dendrite 180 – 200 µm from the soma. The theta-burst pairing (TBP) protocol consisted of a burst of 5 EPSP-current injection pairs delivered at 100 Hz and each burst was delivered 10 times at 5 Hz. This train was repeated three times at 10 s intervals.

### SDS-PAGE and Western blot analysis

Whole hippocampi collected at P60 were used to assess protein expression. Protein concentration was determined using Bio-Rad Protein Assay Dye (Hercules, California, USA; Cat: 5000006). Samples were mixed with SDS sample buffer and 10 μg of protein was loaded in duplicate on SDS-PAGE gels, then transferred to PVDF Transfer Membrane (Millipore Sigma, Darmstadt, Germany). Membranes were blocked using 5% milk for 1 hour. Antibodies were diluted in 1% Tween in PBS or 5% Bovine Serum Albumin (Sigma, CAS # 9048-46-8) prepared in 1% Tween in PBS (filtered) and incubated overnight at 4°C. Membranes were then washed and incubated with secondary antibody, either Rabbit IgG HRP Linked Whole Antibody (Millipore Sigma, Darmstadt, Germany; Cat: GENA934) or Mouse IgG HRP Linked Whole Antibody (Millipore Sigma, Darmstadt, Germany; Cat: NXA931V). Signals were detected with enhanced chemiluminescence using Pierce ECL Blotting Substrate (Thermo Scientific, Carlsbad, CA, USA, Cat:32106) and BioBlue-LiteTM Western Blot Film (Alkali Scientific Inc, Fort Lauderdale, FL, USA, Cat: XR8815). If a second detection was needed, blots were stripped using Restore Western Blot Stripping Buffer (Thermo Scientific, Carlsbad, CA, USA, Cat:21059), blocked again in 5% milk, and incubated overnight with the desired antibody. Blot images are shown in grayscale in the main figures.

Signal intensities of proteins were normalized to GAPDH signal on the same blot. The average of the duplicates was counted as one data point. Protein-specific signals on Western blots were quantified densitometrically using NIH ImageJ software (Bethesda, Maryland, USA).

### RNA isolation and qRT-PCR

RNA was extracted using Trizol® (Life Technologies, Carlsbad, CA). cDNA was generated using High Capacity RNA-to-cDNA Kit (Applied Biosystems, Foster City, CA) for mRNA, or qScript™ microRNA cDNA synthesis kit (Quanta BioSciences, Gaithersburg, MD) for miRNA, followed by SYBR green quantitative real-time PCR (Bio-Rad Laboratories, Hercules, CA). Relative changes were quantified using the comparative cycle threshold method (2−ΔCT). Quality and quantity of mRNA was measured using a Nanodrop Spectrophotometer (Thermo Fisher Scientific, Waltham, MA) or BioTek Cytation Imaging Microplate Reader (BioTek, Winooski, VT) and RNA dilutions were made in nuclease-free water.

Reverse transcription for individual qPCRs was carried out using 500 ng (or maximum volume) of RNA and the High Capacity RNA-to-cDNA Kit (Applied Biosystems, Foster City, CA) using specific primers for miR-324-5p, miR-324-3p, and mRNAs of interest (Quanta BioSciences, Gaithersburg, MD). Individual qPCRs were carried out on the QuantStudio 3 Real-Time PCR System (Applied Biosystems, Foster City, CA) using iTaq Universal SYBR green supermix (Bio-Rad Laboratories, Hercules, CA). A relative fold change in expression of the target gene transcript was determined using the comparative cycle threshold method (2−ΔCT).

### Antibodies, antagomirs, and primers

#### Antibodies

The following antibodies were used: Kv4.2 mouse monoclonal (Antibodies Inc., Cat# 75-016, RRID:AB_1019722), Gli1 rabbit polyclonal (Thermo Fisher Scientific Cat# PA5-17303, RRID:AB_10985784), MAP2 rabbit polyclonal (Millipore Cat# AB5622, RRID:AB_91939), Beta-actin mouse monoclonal (Sigma-Aldrich Cat# A1978, RRID:AB_476692), Alpha-tubulin mouse monoclonal (Sigma-Aldrich Cat# T6074, RRID:AB_477582), GAPDH mouse monoclonal (Abcam Inc Cat# AB9484, RRID:AB_307274), Beta 3 tubulin rabbit polyclonal (BioLegend Cat# 802001, RRID:AB_2564645), Anti-PSD-95 MAGUK scaffold protein mouse monoclonal (Antibodies Incorporated Cat# 75-348, RRID:AB_2315909), Synapsin-1 rabbit polyclonal Sigma-Aldrich Cat# S193, RRID:AB_261457), KCNQ2 mouse monoclonal (Proteintech Cat# 66774-1-Ig, RRID:AB_2882120), and KCNQ3 rabbit polyclonal (Alomone Labs Cat# APC-051, RRID:AB_2040103).

#### Antagomirs

For ICV injections of miR-324-5p-specific or scrambled antagomirs, mice received a 2 μl infusion of 0.5 nmol of either scrambled or miR-324-5p specific antagomirs in aCSF. All antagomirs were locked-nucleic acid-modified and obtained from Exiqon, Vedbaek, Denmark. A custom-made in vivo inhibitor (15 nucleotides) with a partial phosphorothioate backbone and no cholesterol tag (due to problems with synthesis and solubility) specific for miR-324-5p and a scrambled control with the same features were used (both Exiqon).

#### Primers

The following qRT-PCR primers were used: GAPDH for: GGGTTCCTATAAATACGGACTGC; GAPDH rev: CCATTTTGTCTACGGGACGA; Kv4.2 for: GCTTTGAGACACAGCACCAC; Kv4.2 rev: TGTTCATCGACAAACTCATGG; PSD95 for: 5TCTGTGCGAGAGGTAGCAGA; PSD95 rev: AAGCACTCCGTGAACTCCTG; MAP2 for: CTGGACATCAGCCTCACTCA; MAP2 rev:AATAGGTGCCCTGTGACCTG; β-Actin for: ACTGGGACGACATGGAGAAG; β-Actin rev: GGGGTGTTGAAGGTCTCAAA; β-tubulin for: TCGTGGAATGGATCCCCAAC; β-tubulin rev: TCCATCTCGTCCATGCCCT; miR-324-5p: CGCATCCCCTAGGGCATTGGTGT; miR-324-3p: ACTGCCCCAGGTGCTGCTGG.

### Statistics

All analyses were performed by experimenters blinded to genotype, treatment, and sex. Appropriate (parametric or nonparametric) statistical tests (indicated in figure legends) were determined and run using GraphPad Prism version 8 (GraphPad Software, San Francisco, CA). Data with unequal variance was assessed using nonparametric methods. ROUT was used to identify and remove outliers at Q=5%. If outliers were not removed, this is indicated in the figure legends. Sample sizes were determined using R (R Core Team 2020) and published or preliminary results. Significance level was set to α<0.05. All experiments shown are fully powered (power>0.8) unless otherwise indicated. All data points are averaged values for individual mice.

Our data showed no sex effects in Thy1-eGFP^hemi^ or *Mir324* KO mice (Table 1). Both male and female mice were used in each experiment in sex and litter/age-matched pairs.

**Table 1:**
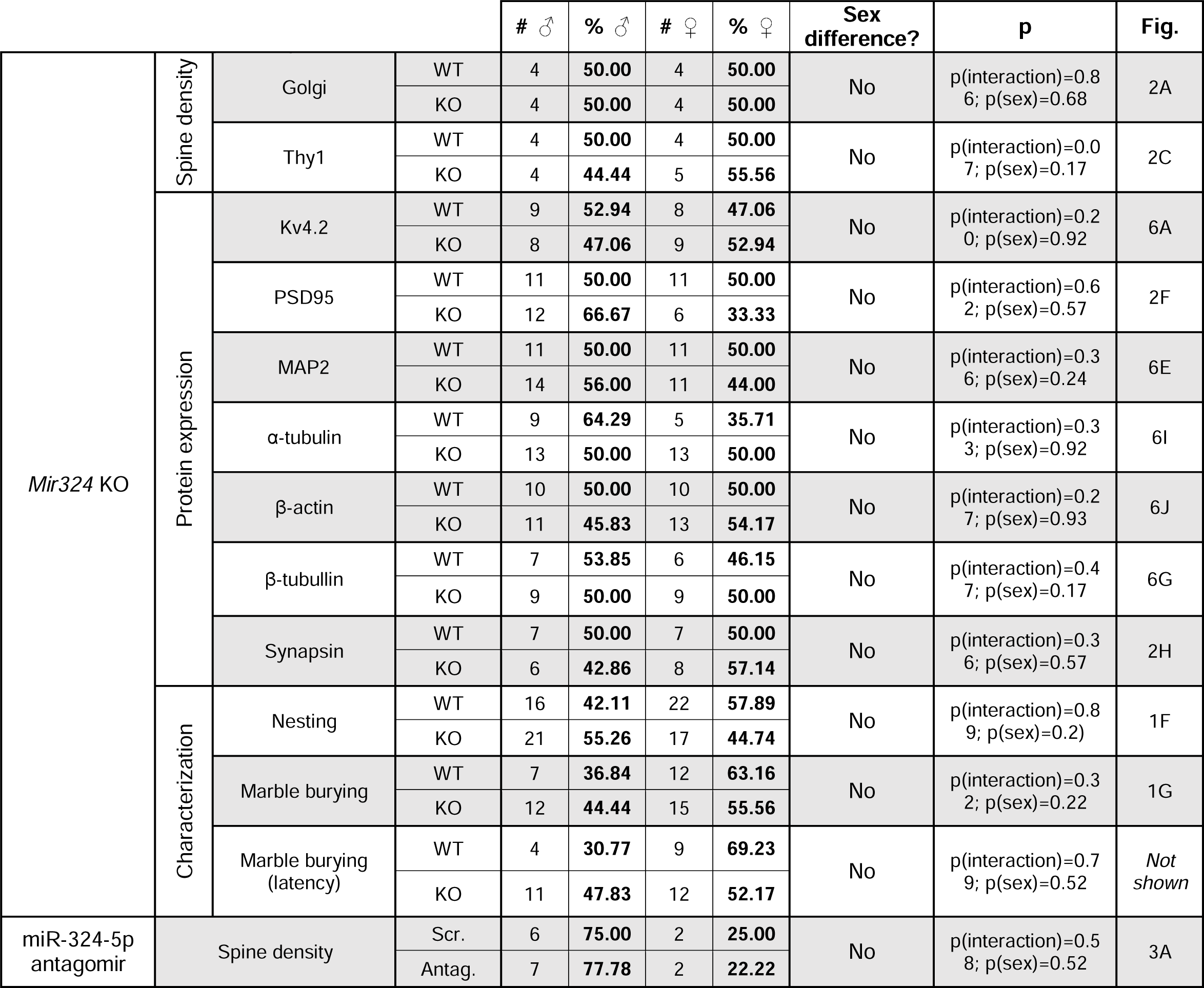
Analysis of sex differences in *Mir324* KO and WT mice. **No sex *differences* were detected in *Mir324* KO and WT mice** (2-Way ANOVA; p>0.05 for all measures). For assessment of spine density and morphology in miR-324-5p or scrambled antagomir treated mice, note that including sex as a variable was underpowered.

## Acknowledgements

This work was funded by National Institutes of Health grants R01NS092705, R01NS107453, R21NS126740, T32NS007453-16, a CCHMC Research Innovation/Pilot Funding Award, and the Cincinnati Children’s Research Foundation. The authors thank the entire CCHMC Veterinary Services team and the CCHMC Confocal Imaging Core for their dedicated support, as well as all members of the Gross lab for thoughtful discussions.

## Data Availability

Data are shown in tables and graphs in the manuscript. Raw data are available from the corresponding author upon reasonable request.

