## Supplemental figures for "MiR-324-5p regulates the structure of dendritic spines and is essential for hippocampal long-term potentiation"

***Correspondence should be addressed to:**

**Content:**

**Supplementary Figure S1**

**Supplementary Figure S2**

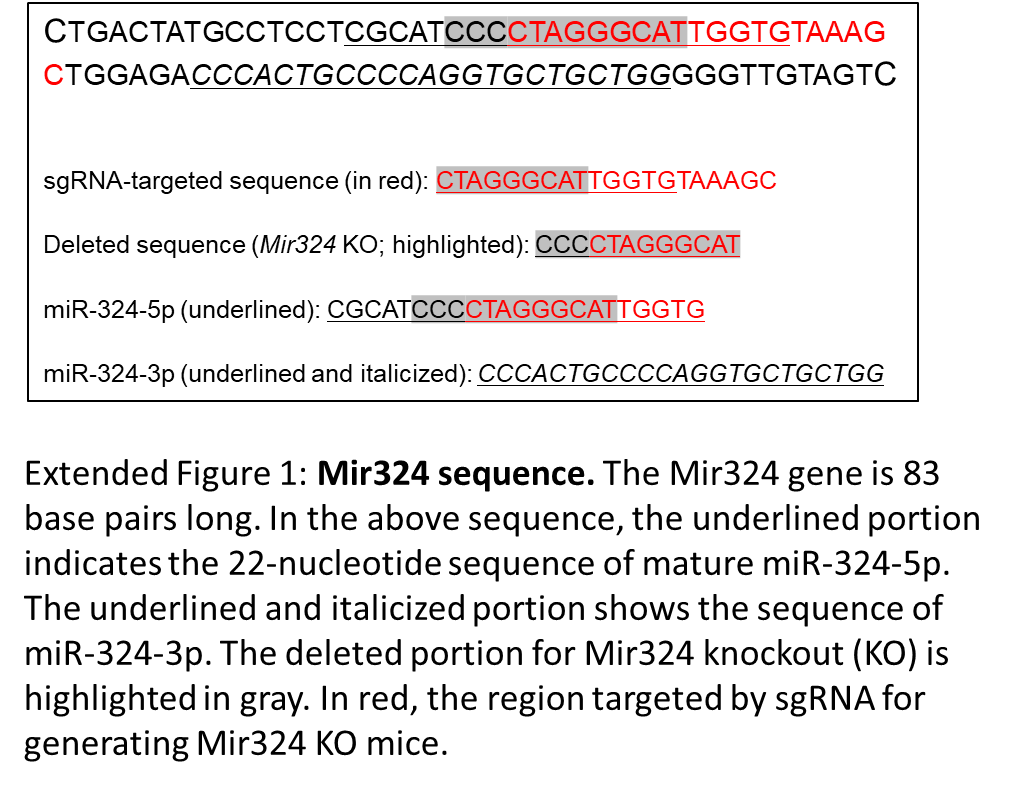
**Supplementary Figure S1**

**Supplementary Figure S1: Mir324 sequence.** The *Mir324* gene is 83 base pairs long. In the above sequence, the underlined portion indicates the 22-nucleotide sequence of mature miR-324-5p. The underlined and italicized portion shows the sequence of miR-324-3p. The deleted portion for *Mir324* knockout (KO) is highlighted in gray. In red, the region targeted by the sgRNA for generating *Mir324* KO mice is shown.

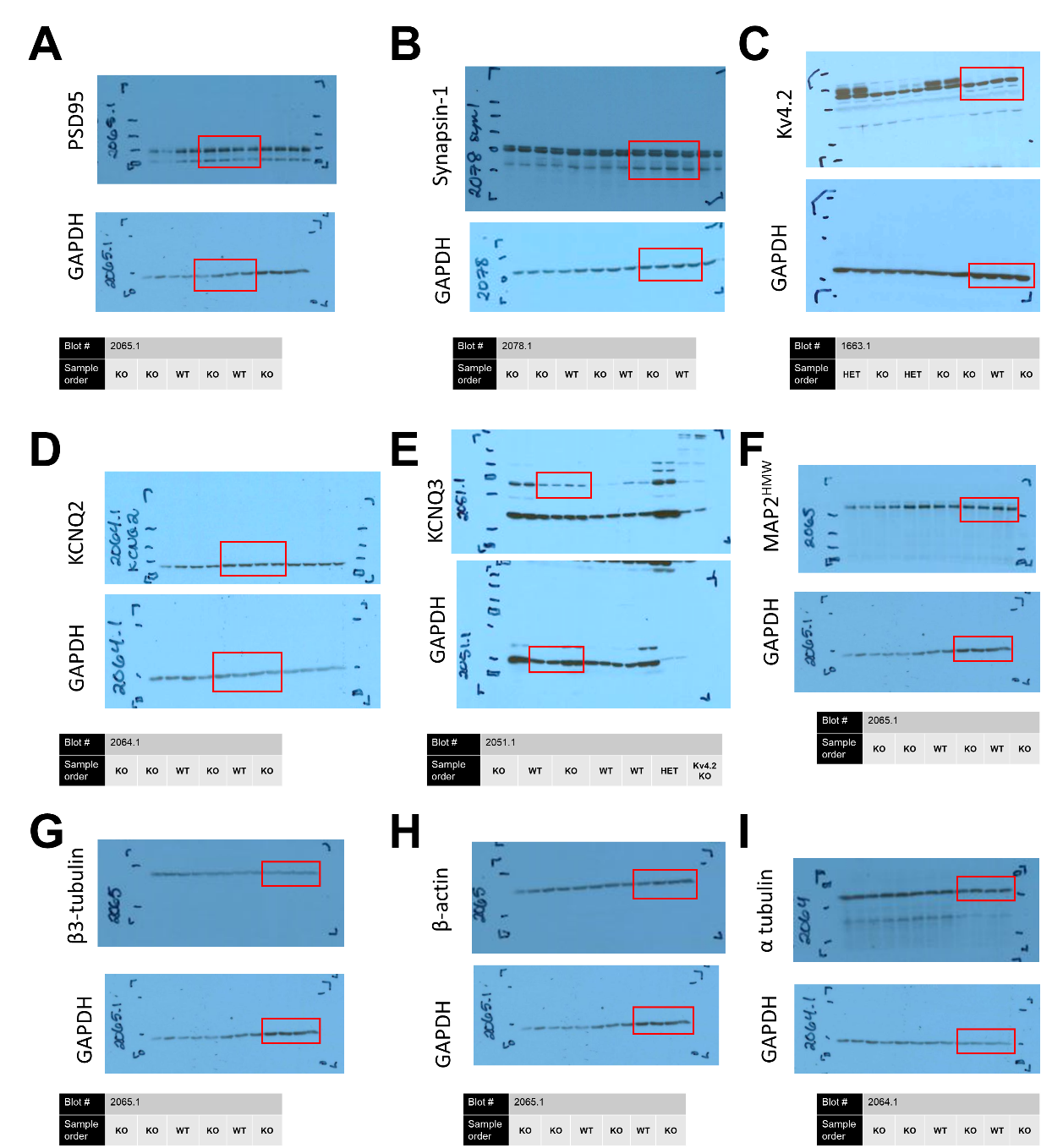
**Supplementary Figure S2**

**Supplementary Figure S2: Western blot images.** Uncropped images of complete western blots shown in Figures 2 and 6. Lanes used for representative images are enclosed in red boxes. The proteins are (**A**) PSD95, (**B**) synapsin-1, (**C**) Kv4.2, (**D**) KCNQ2, (**E**) KCNQ3, (**F**) MAP2^HMW^, (**G**) β3-tubulin, (**H**) β-actin, and (**I**) α-tubulin. Note that several proteins were measured on the same blot. Each hippocampal tissue sample is loaded in duplicate. Blot number and order of genotypes are listed in tables below the blot images.
